# Image-Based Profiling of Induced Trophoblast Stem Cells Identifies Signatures Associated with Sex, Schizophrenia Genomic Risk and Placental Stress

**DOI:** 10.64898/2025.12.23.695254

**Authors:** Frank J. Piscotta, Jiyoung Kim, Jisu Ha, Bonna Sheehan, Julia Johnston, John Peters, Thomas M. Hyde, Brady J. Maher, Juan Caicedo, Daniel R. Weinberger, Evgeny Shlevkov, Gianluca Ursini

**Affiliations:** Lieber Institute for Brain Development, Johns Hopkins Medical Campus, Baltimore, MD, USA; Department of Physiology, Pharmacology and Therapeutics, Johns Hopkins University School of Medicine, Baltimore, MD, USA; Morgridge Institute for Research, Madison, WI, USA; Department of Biostatistics and Biomedical Informatics, University of Wisconsin-Madison, Madison, WI, USA; Department of Psychiatry and Behavioral Sciences, Johns Hopkins University School of Medicine, Baltimore, MD, USA; Department of Neurology, Johns Hopkins University School of Medicine, Baltimore, MD, USA; The Solomon H. Snyder Department of Neuroscience, Johns Hopkins University School of Medicine, Baltimore, MD, USA; McKusick-Nathans Institute, Department of Genetic Medicine, Johns Hopkins University School of Medicine, Baltimore, MD, USA

**Keywords:** Cell Painting, image-based profiling, machine learning, trophoblast stem cells, schizophrenia

## Abstract

Schizophrenia (SCZ) is a neurodevelopmental disorder where both genetic and environmental risks converge during pregnancy. Recent studies have highlighted the importance of placental biology in influencing risk of developing SCZ. However, the pathways by which genetic risk factors for SCZ interact with environmental influences to alter placental development are poorly understood. Through image-based profiling using Cell Painting, we leveraged trophoblast cultures derived from human induced pluripotent stem cells (hiPSCs) from male and female SCZ and neurotypical donors with varying placental genomic risk scores (PlacGRS) to explore the developmental dynamics of placental cells under normal growth and hypoxic stress. We employed both classical (e.g., CellProfiler) and deep learning feature extraction combined with downstream supervised machine learning to analyze high-dimensional data obtained from this hiPSC-derived model system, highlighting that these approaches overcome the inherent line-to-line variability in phenotypic analysis. Our findings reveal a salient nucleus-localized SCZ risk signature across cell lines, along with clear sexual dimorphism. This research underscores the capability of hiPSC-derived placenta models to elucidate complex interactions between genetic risk and environmental factors implicated in the neurodevelopment of SCZ, paving the way for future studies aimed at developing targeted therapeutic and prevention strategies.

## Introduction

Schizophrenia (SCZ) is a heritable polygenic neurodevelopmental disorder affecting approximately 1% of the population worldwide, with higher incidence in males than in females^1,2^. Extensive research has established that risk for SCZ originates, at least in part, *in utero,* whereby genetic risk and environmental factors increase developmental "noise,” which in turn alters neurodevelopmental trajectories^3,4^. Early Life Complications (ELCs)—defined as severe complications during pregnancy, labor/delivery, and the early neonatal period—constitute the most significant environmental risk factors identified for SCZ and other Neurodevelopmental Disorders (NDDs)^5,6^. Specific ELCs that have been associated with SCZ and other NDDs include maternal infection, nutritional deficits, maternal hypertension, preeclampsia, and premature rupture of membranes, as well as outcomes like intrauterine growth restriction, preterm birth, low birthweight, hypoxia, and birth asphyxia^5,6^. Many of these ELCs point to the placenta as a key organ mediating the interaction between genetic and environmental risk in SCZ. ELCs have also been shown to interact synergistically with genomic risk for SCZ, amplifying the liability for the disorder up to fivefold when complications are present compared with when they are not^7,8^.

Mounting evidence supports the hypothesis that many SCZ-associated genomic variants converge on placental mechanisms, leading to altered developmental trajectories that begin in very early life^7–10^. For example, many genes that are associated with SCZ Genome-Wide Association Study (GWAS) risk loci are highly expressed in the placenta, particularly following complicated pregnancies, with expression notably higher in male compared with female offspring^10^. These placental risk loci specifically drive the interaction between SCZ genomic risk, ELCs, and case-control status, as described by the Placental Genomic Risk Score (PlacGRS), a component of genomic risk for SCZ specifically linked to genes highly expressed in placenta. Of note, PlacGRSs are linked with adverse early neurodevelopmental trajectories, as shown by independent associations with neonatal head circumference^9^, neonatal brain volume, and lower developmental scores at one year of age, especially in male individuals^8^. Furthermore, a recent Transcriptome-Wide Association Study (TWAS) and Mendelian Randomization analysis using placental genotype and expression data identified 139 unique and likely causative placental genes associated with SCZ risk^7^. These genes, prioritized by comparison with prenatal cortical brain data, converged on mechanisms related to placental nutrient-sensing and trophoblast invasiveness, such as the inhibition of the mTOR and EIF2 signaling pathways. Among multiple neuropsychiatric illnesses, SCZ exhibited the greatest number of placental genes with a potentially causative role for regulating both expression^10^ and methylation level^11^. These robust and largely replicated findings motivated us to investigate the cellular mechanisms that mediate interactions of genomic risk, sex and environmental stress in placental development in a placenta model system.

Recently, a number of protocols for the differentiation of Trophoblast Stem Cell (TSC)-like cells from human induced pluripotent stem cells (hiPSCs) have been reported ^12–17^, thus permitting the modeling of gene environment interactions in human placenta cells. This capability also enables *in vitro* placental pathophysiology modeling without being restricted by the need for primary placental samples from patients, expanding our ability to investigate cellular mechanisms underpinning interactions between genomic and environmental risk. Human TSC models established from reset naïve-like pluripotent stem cells (PSCs) display self-renewal and maintain stemness at the TSC stage, allowing for maintenance over prolonged periods of time^17–19^. They express the expected genetic and proteomic TSC markers and can differentiate into invasive extravillous trophoblasts (EVTs) and multinucleated syncytiotrophoblasts (STBs), implying that they may mimic physiological placental function *in vitro.* Furthermore, they show matched HLA-class I and C19MC expression and hypomethylation of the *ELF5* promoter region^17,18^. This collective evidence led us to use a naïve iPSC-based protocol for TSC differentiation, which allowed us to generate and maintain large numbers of induced TSC (iTSC) lines for image-based profiling and detection of placental features that are associated with SCZ.

We focus here on iTSC profiling by Cell Painting, an image-based profiling assay ideally suited for studying large numbers of biological samples and perturbations^20,21^. While image-based profiling has been mostly applied to studies of chemical or genetic perturbations in immortalized cancer cell lines^21–23^, some studies have also reported the application of Cell Painting to primary^24,25^ and iPSC-derived cells^26^, demonstrating the application of image-based profiling in pursuit of disease-associated cellular mechanisms. We analyzed Cell Painting images of iTSCs using both CellProfiler^27^ and a pre-trained convolutional neural network (CNN)^28^, and mined the feature and embedding spaces using machine learning (ML)^20^, a strategy that excels at identifying robust signatures in complex scenarios^24,29^. Supervised ML was successful in both sex and diagnosis classification tasks. Leveraging importance values from Random Forest modeling, we extracted key sex and diagnosis-distinguishing features and observed notable differences between male and female lines in terms of those used to distinguish SCZ from control lines. We additionally extracted features selectively affected by hypoxia in male and female lines. We demonstrate that this approach is a fruitful phenotypic exploration of patient-derived lines, opening new avenues for mechanistic investigation and drug discovery in placental health.

## Results

### Research design

Inspired by Schiff et al.’s successful strategy of using Cell Painting and machine learning to map Parkinson’s disease phenotypes from biobanked cell lines^24^, we hypothesized a similar approach could be used to profile hiPSC-derived trophoblast models related to SCZ. We chose 23 fibroblast-derived hiPSC lines from SCZ and neurotypical control (Ctrl) donors, generated and characterized by the Lieber Institute for Brain Development (LIBD), to be differentiated into iTSCs for morphological profiling (**Fig. 1a**). To maximize our chance of establishing a causative link between observed phenotypes and SCZ genomic risk, hiPSCs lines were selected based on PlacGRS, with relatively low PlacGRS in Ctrl and high PlacGRS in SCZ lines (**Table 1, Supplemental Fig. 1a**)^8^. All 23 cell lines were genetically distinct, of European ancestry, and representative of both male and female donors. All female hiPSC lines retained elevated –albeit variable – levels of *XIST*, a marker of X-chromosome inactivation (**Supplemental Fig. 1b**). For sample identity purposes, all fibroblast and hiPSC lines were genotyped and matched to the individuals’ DNA. To generate iTSCs, hiPSCs were reset to naïve-like stem cells, differentiated into induced trophectoderms, and then further differentiated into iTSCs^14,17,30^. iTSC identity was confirmed by immunocytochemistry (ICC) for KRT7, a pan-TSC marker^31^, and the trophectoderm marker GATA3 (**Fig. 1b**)^32^. We further validated our iTSCs by confirming their ability to generate STBs and EVTs. We confirmed the presence of STBs and EVTs by ICC of Syndecan-1 (SDC1), a major transmembrane protein of STB on the placental surface^33^, and human leukocyte antigen G (HLA-G), which is specifically expressed in EVTs and contributes to maternofetal immune tolerance (**Fig. 1c**). The results that follow involve only iTSCs.

**Figure 1:**
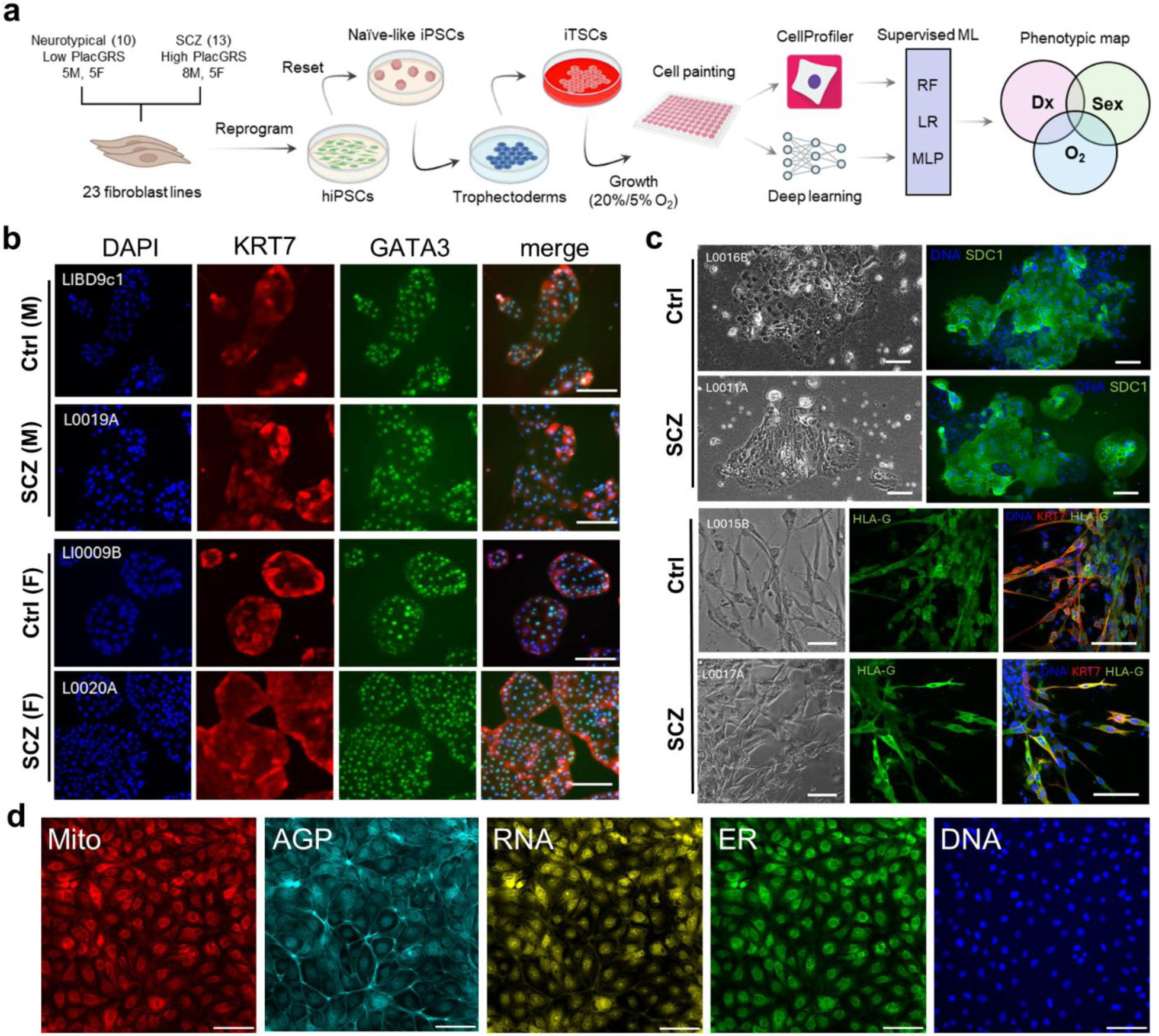
**Study overview and iTSC verification**. **a**) Experimental workflow for Cell Painting characterization of 23 fibroblast lines differentiated into iTSCs. RF: Random Forest, LR: logistic regression, MLP: multilayer perceptron. **b**) Representative immunostaining of male (top) and female (bottom) iTSC lines. Cells were stained with DAPI alongside antibodies for KRT7, a pan-TSC marker and GATA3, a trophectoderm marker. **c**) Representative bright field (left) and immunostained (right) images of iTSCs differentiated into STBs (top) and EVTs (bottom). STB lines 16B (Ctrl) and 11A (SCZ) were stained with DAPI and an antibody for syndecan-1 (SDC1), a transmembrane protein highly expressed on the placental surface. EVT lines 15B (Ctrl) and 17A (SCZ) were stained with DAPI and antibodies for KRT7 and HLA-G, a placental immune marker expressed in EVTs. **d**) Representative images of iTSC line 9B stained with Cell Painting dyes. All scale bars are 100 µm.

**Table 1.** List of cell lines used in this study. PlacGRS: Placental genomic risk score.

| ID (clone name) | Sex | Dx | Age | Fibroblast Source | PlacGRS |
| --- | --- | --- | --- | --- | --- |
| L0001A_X04 | F | SCZ | 32 | Skin | 1.070 |
| L0002A_X03 | F | SCZ | 24 | Skin | 0.7813 |
| L0011A_X05 | F | SCZ | 54 | Brain/Dura | 0.5257 |
| L0017A_X01 | F | SCZ | 18 | Skin | 0.8766 |
| L0020A_X03 | F | SCZ | 51 | Brain/Scalp | 0.9901 |
| L0003B_X01 | F | Ctrl | 27 | Skin | 0.3909 |
| L0004B_X02 | F | Ctrl | 24 | Skin | 0.6287 |
| L0008B_X01 | F | Ctrl | 58 | Brain/Dura | 0.5709 |
| L0009B_X05 | F | Ctrl | 90 | Brain/Dura | 0.5510 |
| L0016B_X01 | F | Ctrl | 23 | Skin | 0.2506 |
| 1002.01 | M | SCZ | 32 | Skin | 1.054 |
| 1006.02 | M | SCZ | 32 | Skin | 1.100 |
| 1013.04 | M | SCZ | 24 | Skin | 1.052 |
| L0006A_Y03 | M | SCZ | 46 | Skin | 0.6659 |
| L0007A_Y03 | M | SCZ | 31 | Skin | 0.9825 |
| L0012A_Y03 | M | SCZ | 28 | Skin | 1.208 |
| L0013A_Y01 | M | SCZ | 40 | Skin | 0.9925 |
| L0019A_04 | M | SCZ | 61 | Brain/Dura | 1.288 |
| 1009.04 | M | Ctrl | 27 | Skin | 0.4081 |
| 1016.02 | M | Ctrl | 39 | Skin | 0.6086 |
| L0015B_Y05 | M | Ctrl | 25 | Skin | 0.4353 |
| LIBD9c1 | M | Ctrl | 52 | Brain/Dura | 0.9114 |
| LIBD7c6 | M | Ctrl | 40 | Brain/Dura | 0.3713 |

Hypoxia represents a common environmental stressor that placental cells experience *in utero* during ELCs^5,6^, and can be modeled *in vitro*. Thus, upon differentiation, all iTSC lines were cultured over two weeks in either a 20% O2 (normoxia) or a 5% O2 (hypoxia) environment. While 20% oxygen is conventionally used for hiPSC-derived TSC culture, we reasoned that experimental switching to 5% oxygen for multiple passages may simulate a pathophysiological hypoxic environment characteristic of pregnancy complications linked with schizophrenia^10,34^. Such a switch is expected to induce chronic hypoxic stress responses and adaptation of cellular physiology to the new environment. We hypothesized that two passages under hypoxia would introduce a broad – yet stable – adaptation of cellular physiology to the new environment, as supported by transcriptomic analyses of the same iTSCs indicating activation of hypoxia signaling genes (**Supplemental Fig. 1c,d**). Following maturation, iTSC lines were seeded in 96-well plates, allowed to attach overnight and underwent Cell Painting using the standard protocol and dye set (DNA, ER, RNA, AGP, Mito) (**Fig. 1e**)^35^.

Images were analyzed using two orthogonal methods – (1) a feature-based approach via CellProfiler (**Fig. 2a**) and (2) a deep learning-based approach (**Fig. 2b**) using a pretrained convolutional neural network (CNN) based on the RepLKnet architecture^28,36^. This allowed us to capture subtle morphological changes using deep learning, while also retaining the biological interpretability associated with traditional features. Approximately 4.3 million cells/crops were analyzed in total. Cell Painting studies, as with other high-content imaging endpoints, can be subject to plate effects, which can complicate downstream analyses by machine learning^37,38^. To help address this issue, we randomized the seeding pattern of cell lines across 12 well-level replicates per line per condition (**Fig. 2c**). We used the DNA-channel focus score, a measure of image quality^38^, as representative metric to examine potential plate effects and did not observe any correlation of focus score to well position (**Fig. 2d, Supplemental Fig. 1d**). Further, for the full dataset, we observed no significant difference in focus scores across columns or rows, indicating no potential bias based on well location (**Fig. 2e,f**). To examine possible batch effects on cell growth, we looked at average cell count per well across plates in the normoxia and hypoxia groups and did not observe significant variation in cell count in the normoxia group (**Fig 2g**). Plates grown under hypoxia did display a slight variation in cell count, suggesting that hypoxia may amplify batch-to-batch variability in growth rate (**Fig. 2h**). Despite this, normalized well-level profiles (both CellProfiler and CNN-derived) across both oxygen conditions displayed no plate-to-plate clustering (**Fig. 2i,j**), mitigating batch effect concerns.

**Figure 2:**
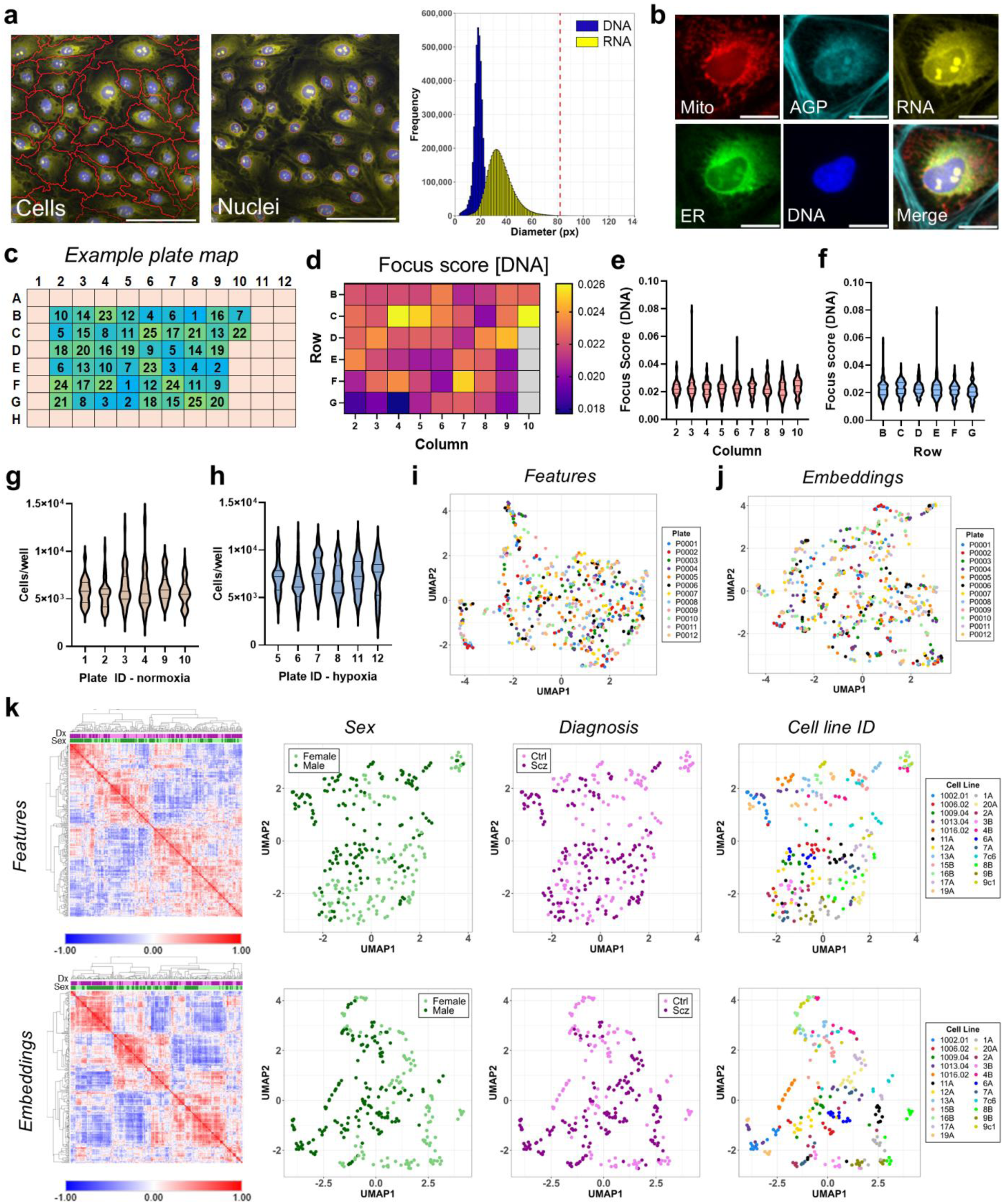
**Experimental setup and quality control**. **a**) Left, example segmentation of cells and nuclei used as input for CellProfiler analysis. DNA (blue) and RNA (yellow) channels used for nucleus and cell segmentation, respectively, are shown. Scale bars = 100 µm. Right, size distribution (in px) of single cell nuclei (DNA) and cell bodies (RNA) across the full dataset. Red dotted line represents the 82 px cutoff used for generating image crops for deep learning. **b**) Representative five-channel image crops (82px x 82px) fed into CNN pipeline. Scale bars = 20 µm. **c**) Randomized plate map for one of the 12 plates used in this study. Each plate contained 25 cell lines (2 wells/line) and seeding was randomized for each plate. Lines 24 and 25 (CT29 and CT30, see methods) were used for assay development but excluded from further analyses as they are not iTSCs. **d**) Well-level focus scores (per-site median, DNA channel) by plate position; normalized median values from across all plates. W = 0.496, p = 0.86 **e**) Violin plots of well-level focus scores from (d) grouped by column. **f**) Same as (e) but grouped by row. W = 2.10, p = 0.066 **g**) Violin plots of well-level cell count grouped by plate ID, normoxia only. W = 1.94, p = 0.093 **h**) Same as (g), but for hypoxia-grown cells. W = 2.87, p = 0.017. Lines on all violin plots mark the median values and the first and third quartiles. All statistical analyses were performed using Welch’s ANOVA. **i**) UMAP of well-level CellProfiler profiles following Pycytominer analysis, grouped by plate ID. **j**) Same as (i) but for deep learning embeddings. **k**) Unsupervised learning of well-level features (top) and embedding (bottom) profiles for normoxia-grown cells. Left, hierarchical clustering (one minus Pearson correlation) of Pearson correlation similarity matrices Entries color coded by sex and diagnosis. Right, UMAPs color coded by cell line sex, diagnosis, and ID.

### Cluster analysis of iTSC lines grown in normoxia

To identify whether cell lines of like sex or diagnosis would cluster under unsupervised learning, we first calculated Pearson correlations for normoxia-only per-well (n=12 wells/line) profiles using either features or embeddings data, followed by hierarchical clustering of the resulting similarity matrix (**Fig. 2k**). While some small clusters of similar sex or diagnosis samples were formed, we did not observe any significant separation of samples along these two classifiers. We then applied a non-linear dimensionality reduction tool (Uniform Manifold Approximation and Projection - UMAP) to the same datasets. When we examined the plots, we still did not observe obvious clustering by diagnosis in either dataset (**Fig. 2l**). When we labeled wells by sex, however, we noted some stratification in the feature space, which was accentuated in the embeddings space (**Fig. 2m**). No clear stratification with these parameters was associated with diagnosis. Finally, we asked whether the sub-clustering of wells in our UMAP plots reflected line to line variance or some other source of variance (such as plate to plate or batch to batch). Consistent with our observations during quality control, the main factor driving sub-clustering was Cell Line ID (**Fig. 2n**). Line to line variance is a common observation in hiPSC studies, which can remain high despite careful experimental quality-controls^39^. Thus, unsupervised clustering indicated that Cell Painted iTSCs show a robust, cell-line specific signature despite cells being grown across six plates and induced in three independent differentiation rounds.

### Identification of features driving sexual dimorphism in iTSCs

Supervised machine learning is a common tool used to analyze Cell Painting datasets (see for example Schiff et al)^24^, while 10-fold cross-validation provides a statistically reliable process to reduce bias in model performance^40^. We constructed a 10-fold cross-validated model for iTSC sex, where each fold contained at least one male and one female iTSC line. (**Fig. 3a and Supplemental Table 1**). Within each fold, we matched, where possible, cell lines by characteristics other than sex such as diagnosis, age of donor and source of the line (skin or dura) to minimize the impact of those variables on the model. Model performance was judged by the area under the curve (AUC) of the receiver operating characteristic (ROC) curve, which measures the ability of the model to correctly rank a random positive sample over a random negative sample^41^. An AUC of 0.5 represents random chance, and model performance improves as AUC approaches 1.0. Because Random Forest (RF) modeling typically has low variance and is relatively robust to overfitting^42^, we trained separate RF models on embeddings and feature datasets. RF performed well at the donor sex classification task (*embeddings* median AUC = 0.88; *features* median AUC = 0.89), suggesting that there are highly salient morphological features distinguishing male and female derived cell lines (**Fig. 3b**). In benchmarking RF performance, comparable AUC values were observed with logistic regression (LR) and multilayer perceptron (MLP) models. Some variation was observed for the embeddings input, where performance was as high as 0.94 median AUC for LR modeling (**Supplemental Fig. 2a**).

**Figure 3:**
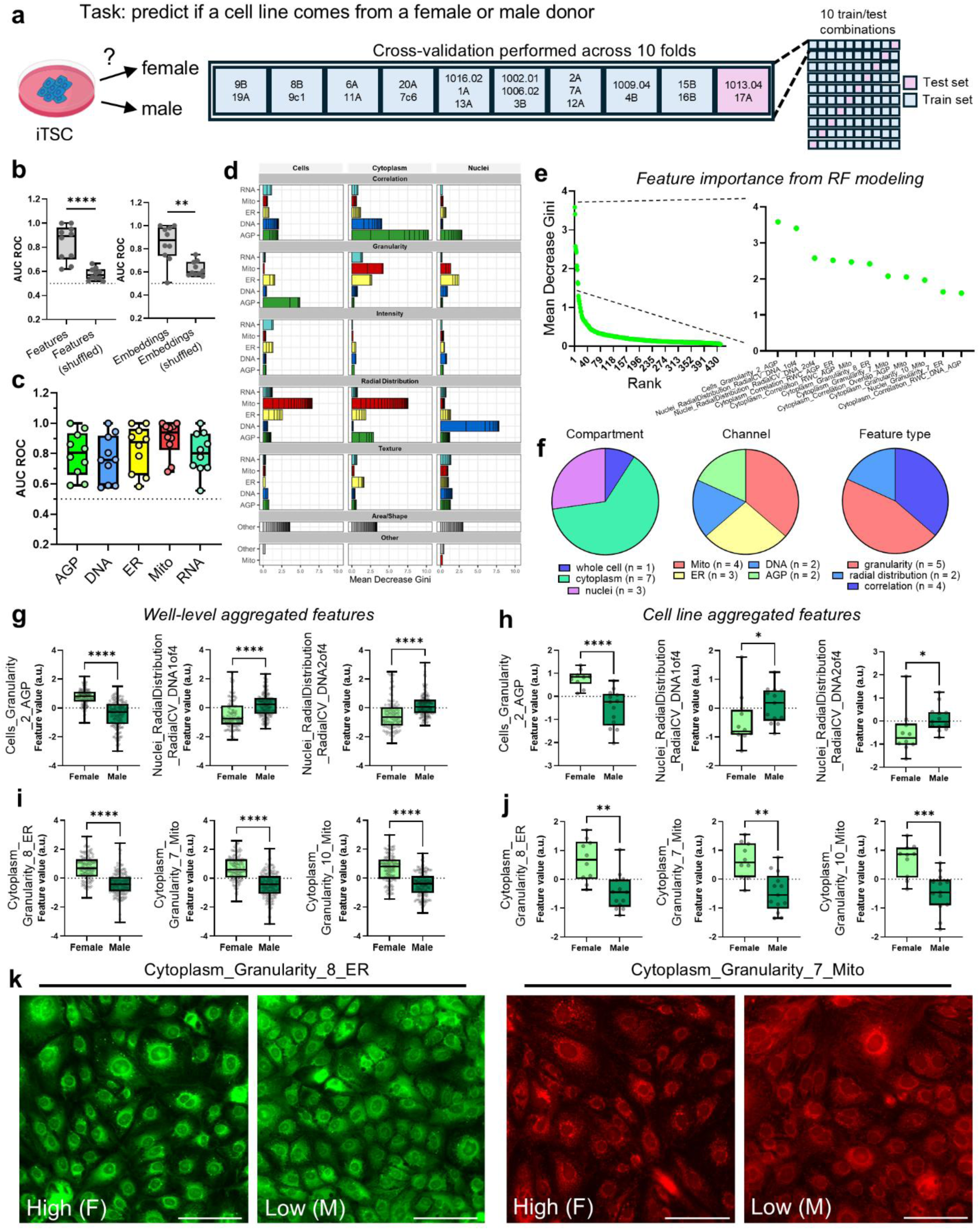
**Sexual dimorphism in iTSC cell lines**. **a**) Schematic overview of Random Forest (RF) modeling with 10-fold cross validation applied to the donor sex classification task. **b**) Area under the curve (AUC) of the receiver operating characteristic (ROC) curve for feature (left) and embedding (right) datasets. Each point represents model AUC for a single testing fold. Performance was compared to the same dataset where male/female labels were randomly assigned. **c**) RF model performance using only single channel embeddings for testing/training. **d**) Representative feature importance plot for the 1016.02-1A-13A testing fold. Features are grouped by compartment, dye channel, and general feature description. Each bar represents a single feature; like features are stacked upon one another. **e**) Feature importance distribution for the testing fold shown in (d) with a focus on the top 11 most important features. **f**) Pie charts showing the compartment, channel, and feature type distributions of the top 11 features. **g**) Well-level comparisons between male and female donor groups of the top three most important features. **h**) Same as (g) but at the cell line level. **i**) Well-level comparisons between male and female donor groups of important cytoplasmic granularity features. **j**) Same as (i) but at the cell line level. **k**) Representative images illustrating the differences in granularity shown in (i) and (j) between female (F) and male (M) lines. Scale bars = 100 µm. Box plots display the interquartile range with a line at the median value. Whiskers extend from minimum to maximum values. All statistical analyses performed using a Mann-Whitney U test (*: p < 0.05; **: p <0.01, ***: p < 0.001, ****: p < 0.0001).

As DINOv3, the latest version of Meta’s self-supervised vision transformer, was recently released^43^, we were curious if analyzing our imaging data using this approach would result in better performance compared to CellProfiler and CNN-based deep learning. We found ML modeling based on DINOv3 outputs to be comparable to our other methods (**Supplemental Fig. 2a**). Of note, though performance did not improve, we obtained strong model performance confirmation in this orthogonal deep learning approach. We were also curious if, in our deep learning output, any single channel embedding was especially critical for classification. We thus performed RF modeling on individual channels rather than the full, concatenated dataset (**Fig. 3c**). Interestingly, RF performance increased when trained purely on the Mito channel (median AUC = 0.94), suggesting that the mitochondrial channel contained sexually dimorphic features.

Next, we looked to the feature-based model to help derive biological insights into the mechanisms driving donor sex-based differences. For each testing fold, we ranked and plotted all 440 features by their importance in performing RF classification using their mean decrease Gini (MDG) values (**Fig. 3d**). Here we present the results of the 1016.02-1A-13A testing fold (AUC = 0.97) though, in general, importance values showed minimal fold-to-fold variation (**Supplemental Table 2)**. We identified 11 features as the strongest contributors to RF classification (**Fig. 3e,f**). When we expanded our analysis of the three most important features to the full dataset, we observed significant differences between female and male groups at the well level (**Fig. 3g**). Since we observed that line-to-line variability drove clustering in unsupervised machine learning, we wondered if this effect was driven by a subset of lines or reflected a generalized trend across all iTSC lines. To test this hypothesis, we calculated the median per-line feature values and performed the statistical test again; significant differences between female and male lines were retained (**Fig. 3h**). Thus, this approach was successful in identifying salient phenotypic differences between groups of iTSC lines.

Of these top three features, *Nuclei_RadialDistribution_RadialCV_DNA_1of4* and *Nuclei_RadialDistribution_RadialCV_DNA_2of4* were significantly increased in male lines. Higher RadialCV in bins one and two indicates greater heterogeneity in DNA staining intensity towards the center of the nucleus, which in biological terms may indicates an uneven distribution of heterochromatin in male iTSCs. Additionally, *Cells_Granularity_2_AGP* was significantly decreased in male lines relative to female lines. Granularity refers to a texture measurement that quantifies the sizes of structures (or "grains") present in an image, with a high value for a small grain size indicating a fine texture (many small features). A lower *Granularity_2_AGP* in male lines suggests increased vesiculation of the Golgi apparatus in female lines compared to male lines. We also noticed that three of the top features were granularity descriptors located in the cytoplasm: *Cytoplasm_Granularity_8_ER, Cytoplasm_Granularity_7_Mito*, and *Cytoplasm_Granularity_10_Mito*. These granularity values correspond to larger elements of texture, and all three saw significant decreases in the male donor population as compared to the female donor population (**Fig. 3i, j**). When we examined the images, we noticed apparent differences in the number and size of granules present in the ER and Mito channels between female and male lines (**Fig. 3k**). Moreover, three cytoplasmic correlation features measuring the coincidence of Mito and ER staining with AGP were significantly increased in male relative to female lines: *Cytoplasm_Correlation_RWC_AGP_ER*, *Cytoplasm_Correlation_RWC_AGP_Mito*, and *Cytoplasm_Correlation_Overlap_AGP_Mito* (**Supplemental Fig. 2b).** Taken together with our deep learning results (**Fig. 3c**), this data points to the existence of sex-dependent differential regulation of ER and mitochondrial dynamics in the cytoplasm of iTSCs.

### Nuclei features are critical for diagnosis classification under normoxia

We then applied the same strategy used for sex classification to identify diagnosis-associated signatures. We performed 10-fold cross-validated RF modeling on the embedding and feature profiles, but this time grouped cell lines so that each fold contained at least one Ctrl and SCZ donor cell line (**Fig. 4a and Supplemental Table 1**). RF models trained on embedding or feature inputs had median AUCs of ∼0.80, indicating a good ability to distinguish between iTSC diagnosis in both datasets. When compared to the same datasets where diagnosis labels have been randomly assigned, both feature and embedding inputs show a statistically significant decrease in RF model performance (median AUCs of 0.56 and 0.60, respectively; P < 0.01), suggesting that this result is not a consequence of overfitting to a large feature space (**Fig. 4b**). LR and MLP models performed similarly, apart from LR modeling performing worse on the embeddings input (**Supplemental Fig. 2c**). As with sex classification, DINOv3 modeling showed similar performance to CellProfiler and CNN-based deep learning. RF models trained using only the DNA or Mito channel embeddings showed similar performance to the full dataset (median AUCs = 0.81, 0.80, respectively), with other channels trending towards lower performance, though the differences were not statistically significant (**Fig. 4c**). This result suggests that iTSCs’ DNA and mitochondria may be important for distinguishing donor diagnosis.

**Figure 4:**
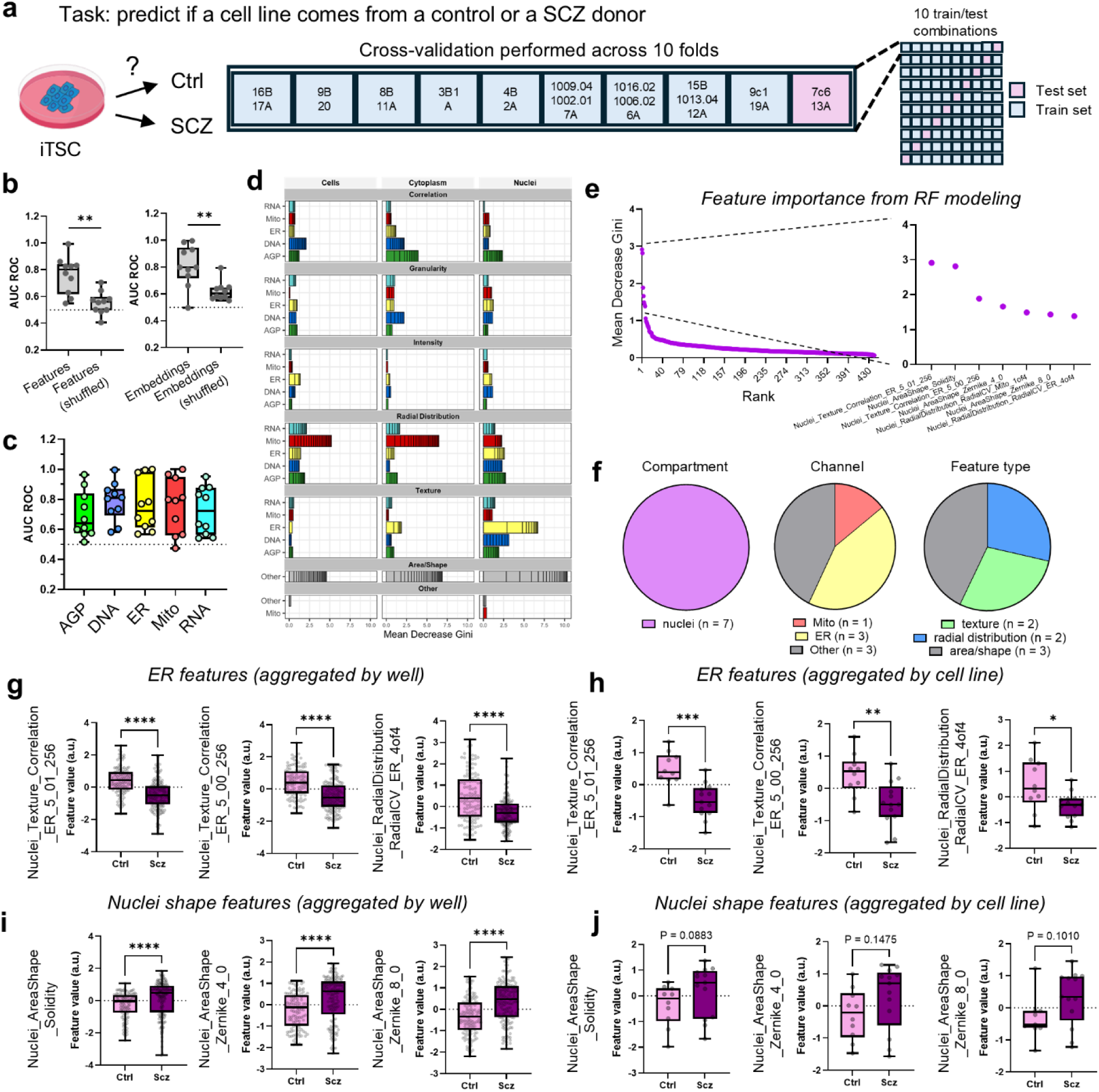
**Nuclear signatures driving diagnosis classification**. **a**) Schematic overview of RF modeling with 10-fold cross validation applied to the diagnosis classification task. **b**) Plots of AUC ROC for feature (left) and embedding (right) datasets. Performance was compared to the same dataset where Ctrl/SCZ labels were randomly assigned. **c**) RF model performance using only single channel embeddings for testing/training. **d**) Representative feature importance plot for the 15B-1013.04-12A testing fold. **e**) Feature importance distribution for the testing fold shown in (d) with a focus on the top seven most important features. **f**) Pie charts showing the compartment, channel, and feature type distributions of the top seven features. **g**) Well-level comparisons between Ctrl and SCZ donor groups of important ER-associated features. **h**) Same as (g) but at the cell line level. **i**) Well-level comparisons between Ctrl and SCZ donor groups of important nuclear shape features. **j**) Same as (i) but at the cell line level. Box plots display the interquartile range with a line at the median value. Whiskers extend from minimum to maximum values. All statistical analyses performed using a Mann-Whitney U test (*: p < 0.05; **: p <0.01, ***: p < 0.001, ****: p < 0.0001).

We again interrogated feature importance in RF modeling (**Fig. 4d and Supplemental Table 3**). Here we focus on the results for the 15B-1013.04-12A testing fold (AUC = 0.99), although, in general, we observed that importance values showed minimal fold-to-fold variation. All top seven features were identified around the nucleus, after which, feature importance dropped substantially (**Fig. 4e**). Of these seven features, three mapped to nucleus area/shape, while the other four were split between ER and Mito signatures in the perinuclear region. Of these four features, two were radial distribution, and two were texture-based (**Fig. 4f**). Thus, nuclei and their immediate environment contain salient phenotypic differences between Ctrl and SCZ iTSC lines in the Cell Painting assay.

Of the top features, two were related to ER texture (*Nuclei_Texture_Correlation_ER_5_01_256* and *Nuclei_Texture_Correlation_ER_5_00_256*) and one to ER distribution in the outermost ring of the nucleus (*Nuclei_RadialDistribution_RadialCV_ER_4of4*). Texture describes the visual roughness or smoothness of an object’s intensity pattern, whereas *Correlation* quantifies the linear dependency of gray levels of neighboring pixels (how similar the intensity of a pixel is to its neighbor). Feature values were significantly decreased in iTSCs derived from SCZ patients compared to controls at the well (**Fig. 4g**) and line level (**Fig. 4h**), indicating a rough, fine, or grainy texture where neighboring pixel intensities vary widely. Since these ER features were identified near the nucleus (i.e., the *Nuclei* feature prefix), decreases in ER correlation and radial coefficient of variation point to irregularities at the nucleus-ER interface. Also of note were three features related to nucleus shape and area (*Nuclei_AreaShape_Solidity*, *Nuclei_AreaShape_Zernike_4_0*, and *Nuclei_AreaShape_Zernike_8_0*). All three feature values were significantly increased in SCZ iTSCs compared to controls at the well level (**Fig. 4i**) and showed numerical trends in the same direction at the cell line level (**Fig. 4j**). Solidity and Zernike are descriptors of shape complexity; simultaneous observations of high solidity alongside high values for the higher-order Zernike moments (e.g., 4 and 8) may describe nucleus shape irregularities in SCZ iTSCs, potentially linked with impaired syncytialization, or pathological differentiation states. One possible underlying mechanism is nuclear envelope (NE) integrity loss and fragmentation. The NE is contiguous with the ER and is a key determinant of chromatin organization^44^. When the NE/ER system breaks down, the organized chromatin structure becomes disorganized or dispersed, which can lead to lower ER feature values and larger Nuclei Zernike and solidity values described above. Taken together, our model suggests mechanisms regulating the NE could be a key feature of illness in this cellular system.

### Hypoxia induces changes in diagnosis signatures in iTSCs

We reasoned that hypoxia-associated changes in the features deemed important for sex or diagnosis classification may point to the cellular substrates of synergies between diagnosis, sex, and placental stress. As with cells grown under normoxia, we first performed unsupervised learning to look for iTSC line clustering by sex or diagnosis (**Fig. 5a**). We did not observe obvious clustering by sex or diagnosis, though wells still tended to cluster by cell line ID. ML models were again successful in classification by sex (**Fig. 5b**). For sex classification, when compared to normoxia, modeling under hypoxia exhibited comparable performance (*embeddings* median AUChyp = 0.89 vs AUCnorm = 0.88; *features* median AUChyp = 0.84 vs AUCnorm = 0.89). LR and MLP models slightly outperformed RF hypoxia modeling (**Supplemental Fig. 3a**). AGP and Mito channels displayed the most discriminatory power in single-embedding tests (AGP AUC = 0.88; Mito AUC = 0.97). As under normoxia, ML classifiers consistently detected sexually dimorphic traits in iTSCs grown under hypoxia, indicating constitutive sexual dimorphism in organelle networks that withstand placental stress. By contrast, diagnosis classification under hypoxia exhibited decreased performance for the embedding dataset (AUChyp = 0.69 vs AUCnorm = 0.80), and comparable feature performance (AUChyp = 0.75 vs AUCnorm = 0.80) (**Fig. 5c**). LR and MLP models also showed generally poor performance (**Supplemental Fig. 3b**). Single channel modeling reveals that DNA and Mito channels, which had shown good classifier performance under normoxia, show performance decreases mirroring overall model performance (AUCDNA_norm = 0.81 vs AUCDNA_hyp = 0.73 and AUCMito_norm = 0.80 vs AUCMito_hyp = 0.69). Information stored in these channels, which had previously driven deep learning-based diagnosis classification, can no longer distinguish between Ctrl and SCZ iTSC. These observations suggest that, in contrast to sex, some of the important features driving diagnosis classification under normoxia might be obfuscated by hypoxic growth.

**Figure 5:**
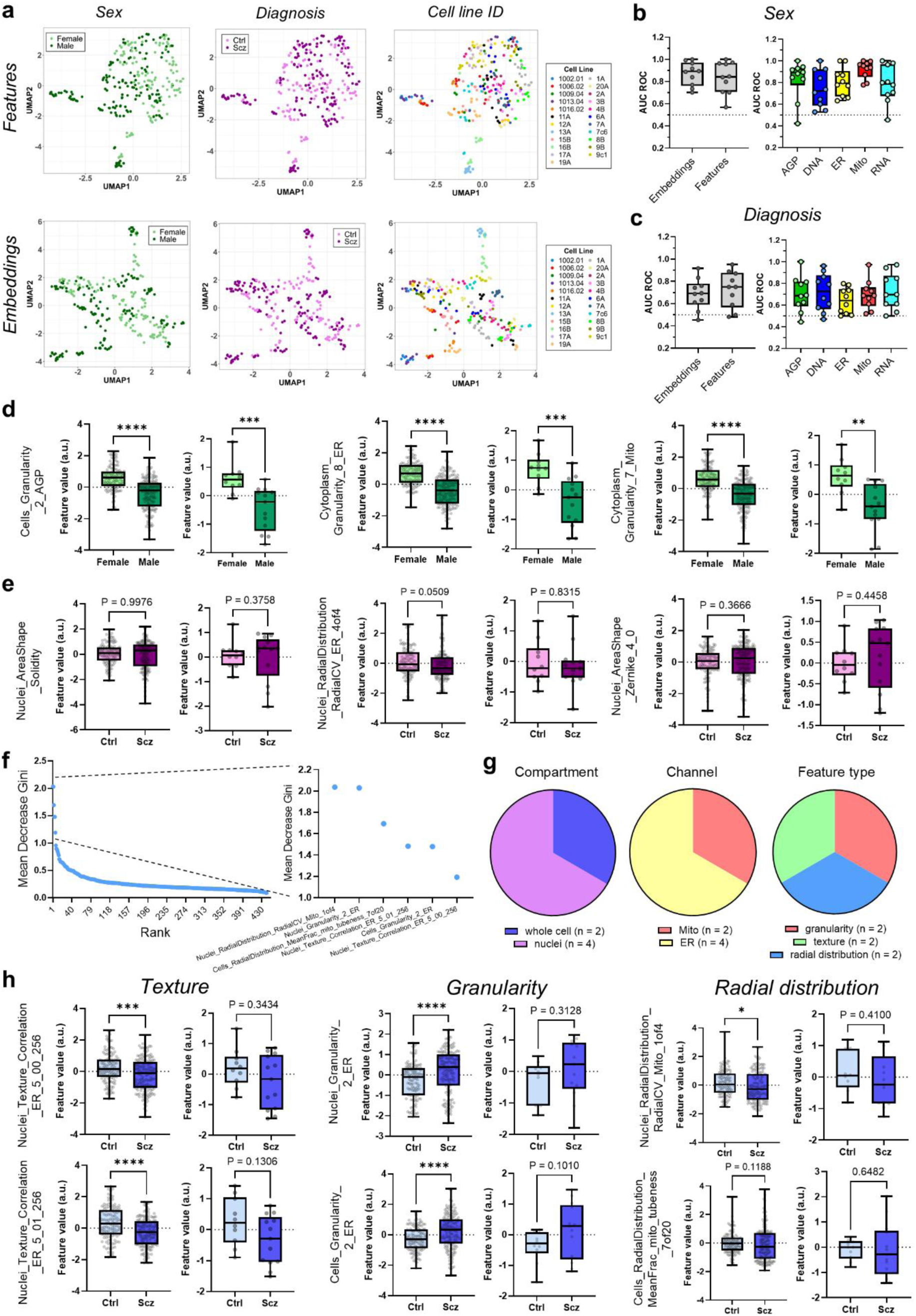
**Hypoxia selectively influences diagnosis classification modeling**. **a**) UMAPs of well-level feature (top) and embedding (bottom) profiles for hypoxia-grown cells. UMAPs are color coded by cell line sex, diagnosis, and ID. **b**) Left, AUC ROC plot comparing feature and embedding model performance under hypoxia. Right, RF model performance using only single channel embeddings for testing/training. **c**) Same as (b) but for diagnosis classification under hypoxia. **d**) Well-level comparisons of male and female groups grown under hypoxia for select features that were important for sex classification under normoxia. **e**) Same as (d) but for Ctrl and SCZ groups in diagnosis classification. **f**) Feature importance distribution for diagnosis classification under hypoxia of the 15B-1013.04-12A testing fold, with a focus on the top six most important features. **g**) Pie charts showing the compartment, channel, and feature type distributions of the top six features. **h**) Well-level and cell line-level comparisons of Ctrl and SCZ groups for top features identified in (f), grouped by general feature descriptors. Box plots display the interquartile range with a line at the median value. Whiskers extend from minimum to maximum values. All statistical analyses performed using a Mann-Whitney U test (*: p < 0.05; **: p <0.01, ***: p < 0.001, ****: p < 0.0001).

Next, we investigated whether hypoxia affected the distributions of the most important features driving sex and diagnosis classification under normoxia. Overall trends in the magnitude and direction of feature value differences between female and male lines identified under normoxia remained high in iTSCs grown under hypoxia (**Fig. 5d and Supplemental Fig. 3c**). However, when we turned to diagnosis, we observed a general trend towards loss of feature salience (**Fig. 5e**). While features describing nuclear ER texture remained significantly different between Ctrl and SCZ groups at the well level, they lost significance at the line level (**Supplemental Fig. 3d**). In total, we observed a pronounced erosion of morphological differences in nuclear shape between Ctrl and SCZ iTSC lines under hypoxia.

We wondered, then, if new features may be driving iTSC line classification under hypoxia. We analyzed the feature importance profile of 15B-1013.04-12A fold (median AUChyp = 0.81), and six features stood out (**Fig. 5f, g**). In contrast to normoxia (**Fig. 2e**), nuclei area/shape features were not driving the classification task. When we plotted individual feature values (**Fig. 5h**), five out of the six features were significantly different only at the well level, indicating large line to line variance in this feature space. We noted that the most significant features corresponded again to the ER channel, whereas the features of lower significance were obtained from the Mito channel. As under normoxia, *Nuclei_Texture_Correlation_ER_5_00_256* and *Nuclei_Texture_Correlation_ER_01_256*, were significantly decreased at the well level in the SCZ lines, indicating that the biological mechanisms driving their importance are resistant to erosion under hypoxia. Two of the four new features corresponded to ER granularity (*Nuclei_Granularity_2_ER* and *Cells_Granularity_2_ER*). In both cases, granularity was increased in SCZ lines compared to Ctrl at the well level, with numerical trends at the line level. Finally, two features suggested potential differences in mitochondrial radial distribution: *Cells_RadialDistribution_RadialCV_Mito_1of4* and *Cells_RadialDistribution_MeanFrac_mito_tubeness_7of20*. However, despite their importance in this particular fold, the features showed little difference between diagnosis groups, suggesting that they are not representative of the general trends in the data. Overall, these data indicate that exposure to hypoxia increases the similarity between controls and SCZ iTSCs in terms of nuclear shape complexity, while greater ER granularity in SCZ iTSCs may indicate coarse ER tubule clustering under stress.

### Sex-specific features in diagnosis groups

Intrigued by the disparity between line-level and well-level significance in certain diagnosis-relevant features, we next tested if the line-to-line heterogeneity was driven by sex. We grouped the salient features (in normoxia and hypoxia) in five classes: *texture_ER*, *shape_nuclei*, *granularity_ER*, *radial_distribution_ER* and *radial_distribution_Mito*, binned the lines by sex and tested for significant feature differences once again (**Fig. 6a**). We observed that, in general, female lines showed significant differences in feature values between Ctrl and SCZ groups across nearly all feature classes. Male group differences reached significance only in one feature: *Nuclei_Texture_Correlation_ER_5_01_256*, and only under normoxia. Thus, we confirmed our hypothesis of the sex-dependence of features critical to diagnosis classification and discovered that the bulk of the difference is driven by female lines. While this observation can be partially attributed to a larger line to line variation between male lines as compared to female lines, our data suggest that an interaction between sex and SCZ risk can be detected in the Cell Painting assay in iTSCs.

**Figure 6:**
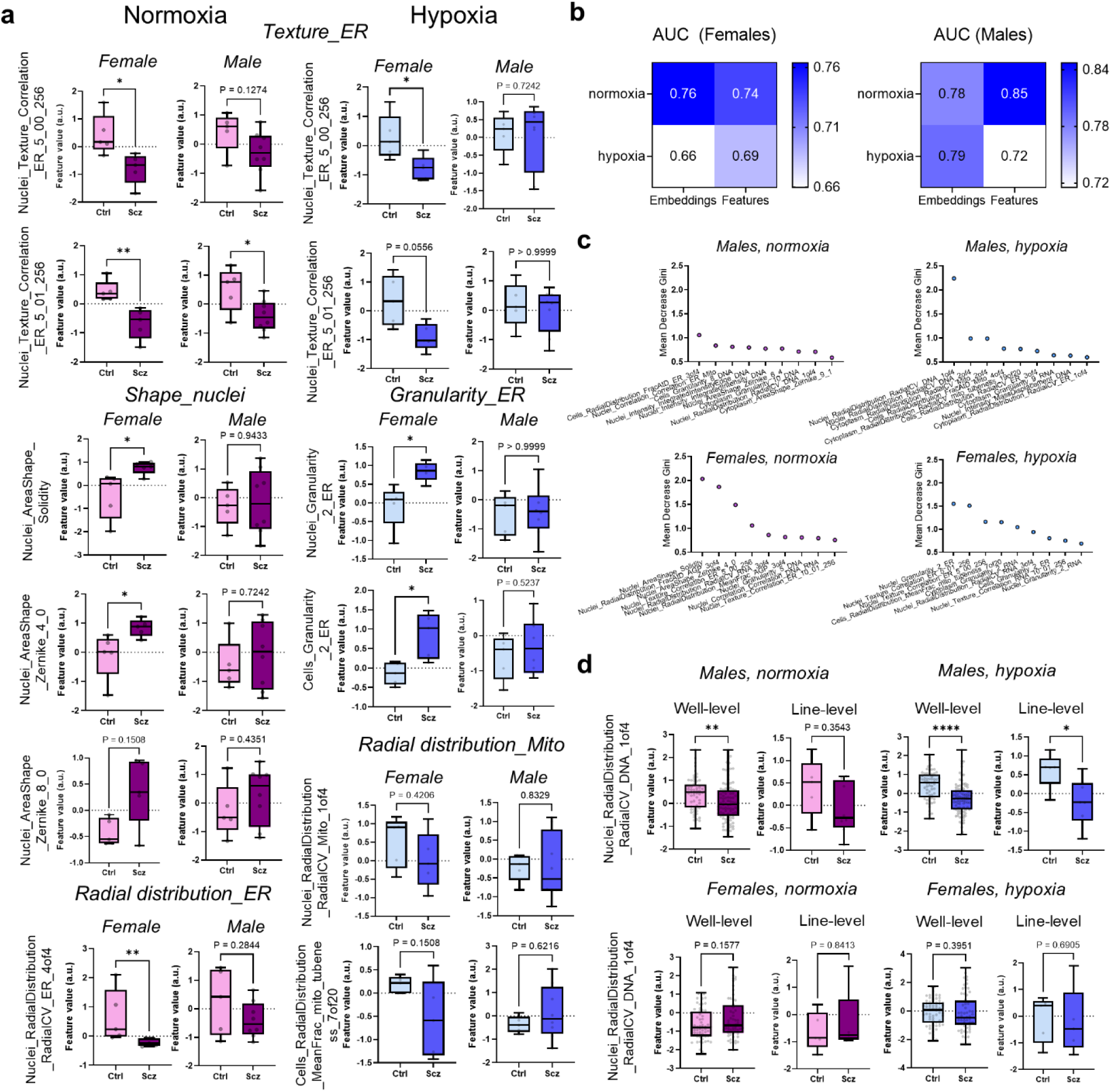
Important diagnosis features display differential behavior in male and female donor lines**. a**) Cell line-level comparison of high importance features for diagnosis classification for cells grown under normoxia (left) or hypoxia (right). AUCs have been separated into male-only and female-only folds. Features are grouped by general descriptor terms. **b**) Matrices depicting RF model performance under various conditions when cell lines are divided by donor sex prior to model training/testing. **c**) Top nine most important features for RF modeling carried out in (b). Plots are split by donor sex as well as oxygen condition during growth. **d**) Well-level and cell line-level comparisons of Ctrl and SCZ groups for feature *Nuclei_RadialDistribution_RadialCV_DNA_1of4,* under normoxia and hypoxia, split between male and female groups. Box plots display the interquartile range with a line at the median value. Whiskers extend from minimum to maximum values. All statistical analyses performed using a Mann-Whitney U test (*: p < 0.05; **: p <0.01, ***: p < 0.001, ****: p < 0.0001).

As epidemiological and gene expression data points to higher SCZ risk potentially linked with the male placenta^7,10^, we were surprised that phenotypic differences between Ctrl and SCZ groups was driven mainly by female lines. Thus, we decided to re-run our diagnosis classification models, this time separating the lines by sex before training, with the objective of identifying new sex-specific signatures. Sex-specific models showed variable performance, with male models generally outperforming females (**Fig. 6b, Supplemental Fig. 4a,b**). We then mined the most important features in male and female datasets (**Fig. 6c**). Several features were consistently among the most important in all five folds in male cell line modeling, with *Nuclei_RadiaDistribution_RadialCV_DNA_1of*4 clearly of top importance under hypoxic conditions. This feature, possibly indicative of an altered chromatin response to hypoxia, was significantly reduced in SCZ male lines compared to Ctrl lines at the well level and reached statistical significance at the line-level under hypoxia; no significant differences were observed in female lines (**Fig. 6d**). Interestingly, this feature had also been identified as a key differentiator in sex classification modeling, so its appearance in diagnosis modeling only when separating lines by sex and its significance uniquely in male lines further accentuates the relevance of sexual dimorphism towards disease-relevant features. Additionally, its increased significance under hypoxia may suggest synergy with the perturbation, whereby hypoxia accentuates the latent interactions between genomic risk and sex. These results appear to echo findings in placenta tissue that show greater molecular effects of genetic risk for schizophrenia in placentae from male offspring compared with female offspring^8,10^.

## Discussion

The nature of the interaction between genomic risk for neurodevelopmental disorders and environmental factors during development remains unclear. HiPSC-derived trophoblast cells derived from patients at opposite ends of the Schizophrenia Genomic Risk Score spectrum offer an experimental platform to investigate such interactions. In this study, we employed image-based profiling by Cell Painting followed by supervised machine learning to examine the cellular characteristics underlying sex differences and SCZ diagnosis in iTSCs under normoxia and hypoxia. Our findings demonstrated a pronounced sexual dimorphism among iTSCs. Notably, nuclear features, specifically those related to area and shape, as well as ER features around the nucleus, were critical in distinguishing between diagnosis groups. We observed changes in the feature valence driving diagnosis but not sex classification upon hypoxia, implicating cellular features at the intersection of SCZ risk and placental stress. These results provide insights into the phenotypic divergence associated with SCZ risk at least in iTSCs, emphasizing the necessity for a more nuanced understanding of sex-specific effects on placental development. Moreover, they demonstrate an analytical framework with the potential to deliver novel mechanistic insights into disease and offer a platform for phenotypic drug screening in patient-derived cells.

One longstanding question in the field is how to extract robust phenotypic information from hiPSC lines in the face of variability and low number of genotypes. Indeed, in unsupervised ML, we observed line-to-line clustering consistent with a large body of evidence, mainly derived from molecular profiling assays such RNA sequencing, indicating that hiPSC-derived lines tend to occupy their own phenotypic space^45–47^. Given that batch effects were not a major factor of variation, and the main clustering was driven by variable cell lines and other demographic characteristics, we reasoned that supervised machine learning offered better control over the complexity of the dataset. To mitigate the potential impact of overfitting, we have 1) employed ten-fold cross-validation, which provides a less biased measure of model performance, 2) analyzed the predictions of three ML methods and 3) leveraged transfer learning from RepLKnet and DINOv3 to constrain our model in deep learning analyses. Our orthogonal strategy proved effective in capturing the subtle complexities of our high-dimensional data sets, showing robust classification performance by ROC AUC (**Fig. 3b, 4b**). We propose that supervised learning can overcome some of the limitations of unsupervised learning on datasets derived from hiPSC lines in part because supervision helps to identify features that can explain the target type of variation such as sex and other demographic characteristics. We believe our framework can be generalized to other studies aiming at phenotypic profiling of patient-derived cell models.

As suggested before^24^, the application of RF importance plots presents a viable way to enrich image-based profiling with mechanistic insights. While biological interpretation of CellProfiler features is not always straightforward, feature analysis may provide testable hypotheses guiding mechanistic investigations. When classifying iTSC lines grown under normoxia by sex, we identified distinct morphological features crucial to differentiating male and female cell lines (**Fig. 3g-j**). Two *DNA Radial Distribution* features suggest differences in heterogeneity of the spatial organization of chromatin. Chromatin heterogeneity is linked to gene expression: transcriptionally silent domains reside at the nuclear periphery, whereas active domains locate within the interior^44^. Chromatin organization differences may underlie sex-biased gene regulation and trophoblast differentiation programs that contribute to the differences in placental development and function^48,49^ and pregnancy outcomes^49–51^ between males and females. Additionally, several important granularity features were also identified in our analysis. We observed one AGP granularity feature was significantly lowered in male cell lines, which is consistent with male placentae having enhanced trophoblast invasion/migration via a stabilized cytoskeleton, and females having finer Golgi fragmentation linked to secretory stress responses (e.g., higher hCG packaging)^49^. Three additional granularity features suggested sex-dependent differences in ER and mitochondrial dynamics. Significant granularity decreases in male cell lines may indicate tighter organelle-plasma membrane apposition in male trophoblasts, which may prioritize mitochondrial fusion/biogenesis for energy-intensive growth. These sex-dependent variations in ER and mitochondria likely reflect distinct metabolic and stress adaptation pathways, highlighting the importance of considering fetal sex in placental disease models and therapeutic strategies.

Separately, feature importance analysis of diagnosis classification revealed nuclear morphology and ER texture as dominant signatures distinguishing SCZ from control iTSCs, implicating the nuclear envelope (NE)–ER–chromatin axis as a convergent locus of disease risk. When nuclear envelope and lamina integrity are compromised, NE and inner nuclear membrane proteins may redistribute into the ER and the perinuclear ER network may become disorganized. In parallel, chromatin that is normally anchored at the NE can lose its ordered peripheral arrangement and become repositioned or compacted, reflecting disrupted higher-order chromatin organization. Such remodeling is expected to reduce ER texture and radial distribution feature values at the nuclear periphery, consistent with the lower ER feature values described above^52,53^. NE/ER interface deviations suggest a mechanism where SCZ risk disrupts perinuclear ER cisternae continuity essential for Ca²⁺ buffering, UPR activation, and lamin-mediated chromatin tethering, complementing transcriptomic evidence that SCZ placental risk genes^7,8,10^ enrich for ER stress/UPR pathways in trophoblasts. Concurrently, elevated nuclear solidity along with high Zernike moments indicate irregular contours, consistent with NE blebbing, lamin dysregulation, or impaired syncytialization, as LMNA remodeling governs nuclear deformation during TSC-STB transition. These structural phenotypes provide orthogonal, cell-autonomous validation of SCZ placental liability, demonstrating how interpretable image-based profiling uncovers subcellular mechanisms bridging genomic risk to trophoblast maladaptation^24^.

In hypoxic conditions, the characteristics of sex-distinguishing features remained relatively stable, whereas the diagnosis-associated features exhibited notable alterations (**Fig. 5d,e**). For instance, while some features mapping to the ER channel continued to significantly differentiate Ctrl and SCZ lines at the well level, even under low oxygen tension, features describing nuclear shape lost their statistical significance, suggesting that hypoxia might dull these distinguishing characteristics. It is worth noting, though, that we have observed that cell counts were more variable under hypoxia as compared to normoxia (**Fig. 2h**), which may have occluded nuclear shape descriptors. However, hypoxia-only features also emerged and suggested synergistic interactions between hypoxia and diagnosis among several ER features. One interpretation could be that these features represent failed tubule elongation and perinuclear retraction, hallmarks of maladaptive UPR, in which SCZ trophoblasts accumulate dilated ER sheets rather than dynamic networks^54,55^. Future studies leveraging image-based profiling and pharmacological tools selectively targeting ER stress^56^ could address this hypothesis.

Important diagnosis classification features identified under hypoxia showed no significance when making line level comparisons, prompting us to examine the effects of donor line sex. This revealed striking disparities between female and male iTSC lines in the context of SCZ diagnosis This sex-specific disparity suggests that genetic and biological frameworks influencing SCZ risk may operate differently in males versus females, highlighting the importance of incorporating sex as a biological variable in both research design and interpretation of results. Our analysis showed that female iTSC lines generally exhibited more significant differences in feature values between Ctrl and SCZ groups across various classes of features

(**Fig. 6**). This disparity suggests that female lines drive most of the observed phenotypic differences associated with SCZ, while male lines derived from SCZ patients showed less consistent salience of features, though often showing similar trends. Since our previous data indicated heightened vulnerability of male placentas to SCZ risk and ELC^8,10^, we were particularly interested in male-specific features. Upon reanalysis, where only male lines were used for testing and training, we found that *Nuclei_RadiaDistribution_RadialCV_DNA_1of4* was one of the few features changing with diagnosis only in males, with the change accentuated under hypoxia (**Fig. 6c**). This implicates an altered epigenetic response to hypoxia in male SCZ trophoblasts.

Our study has some limitations. iTSCs are an undifferentiated precursor cell type, and thus the effects of the interactions of sex, diagnosis and hypoxia need to be further analyzed in differentiated cell types such as STBs and EVTs. These cell types may better approximate the biological milieu of the placenta. We are also limited by the low *n* of genotypes, and thus, whether our findings are generalizable to a larger sample remains to be determined. Further, hypoxia treatment introduced a noticeable increase in variability in cell proliferation, which may have influenced the overall feature valence. Nevertheless, our study demonstrates that image-based profiling with Cell Painting of hiPSC derived cellular models of trophoblasts uncovered sex and diagnosis-associated phenotypic differences. Overall, these findings reveal a significant sex-by-diagnosis interaction shaping cell morphological traits, indicating that genomic and environmental factors differentially influence male and female lines in the context of schizophrenia risk via placenta mechanisms.

## Methods

### Generation of iTSCs, SBs and EVTs

Reprogrammed hiPSCs were selected from the LIBD hiPSC repository generated as described earlier^57,58^. Briefly, primary fibroblasts were collected from dura mater from postmortem samples, part of the LIBD brain repository, and skin biopsies from living subjects from the Sibling Study of Schizophrenia at the National Institute of Mental Health in the Clinical Brain Disorders Branch (NIMH, Protocol 95-M-0150, NCT00001486, Annual Report no. ZIA MH002942053, DRW., PI) with additional support from the Clinical Translational Neuroscience Branch, NIMH (Karen F Berman, PI). Fibroblasts were cultured in fibroblast medium consisting of DMEM with GlutaMAX, 10% fetal bovine serum and 1x Antibiotic-Antimycotic at 37°C, 5% CO2. Fibroblasts underwent Sendai virus reprogramming^57^ or episomal vectors reprogramming^58^. After selection of colonies, hiPSCs were moved to feeder-free culture on 1% Matrigel (Corning, cat. 356230) pre-coated plates in StemFlex^TM^ medium (Thermofisher Scientific, cat. A3349401) at 37°C, 5% CO2, and 20% O2 with humidity, which is called normoxia hereafter. Selected hiPSCs were qualified for their stemness and differential potential following the ISSCR guidelines as well as confirmed Short Tandem Repeat (STR) matching with original donor fibroblasts. Qualified feeder-free hiPSCs between passage 9-12 were then reset to naïve-like state in RSeT^®^ feeder-free (RSeT-FF) medium (Stem Cell Technologies, cat. 5975) supplemented with 0.2% Matrigel (Corning, cat. 354277) following the manufacturer’s protocol. Naïve-like cultures were kept at 37°C, 5% CO2, and 5% O2 with humidity, which is called hypoxia hereafter. Naïve-like PSCs within passage three were differentiated. Reset naïve-like PSCs were plated on 5 μg/ml Collagen IV pre-coated plates in hypoxia a day before, then fed trophectoderm (TE) medium for 2-3 days keeping the culture in hypoxic condition. TE medium was adapted from Guo and colleagues^30^ as follows; 1:1 DMEM/F12 with GlutaMAX and neurobasal, 1x B27, 1x N2, 0.1 mM β-mercaptoethanol, 1 µM PD0325901, and 1 µM A83-01^30^. We switched oxygen concentration from 5% to 20% O2 to introduce the TS medium, which is composed of the DMEM/F12 with GlutaMAX supplemented with 0.1 mM β-mercaptoethanol, 0.2% fetal bovine serum, 0.3% bovine serum albumin, 1% ITS-X, 1.5 ug/ml L-ascorbic acid, 50 ng/ml EGF, 2 µM CHIR99021, 1 µM SB431542, 0.8 mM VPA, 5 µM Y27632, and 1 x Antibiotic-Antimycotic (Thermofisher Scientific, cat. 15240062)^14^. The iTSCs were kept in normoxia and the medium was refreshed every day until the first passage where they grew to 80-90% confluency. iTSCs were confirmed by anti-KRT7 (Thermofisher Scientific, cat. MA5-11986) and anti-GATA3 (Thermofisher Scientific, cat. PA1-101) immunostaining. Differential potential into syncytial trophoblasts (STBs) and extravillous trophoblasts (EVTs) was confirmed by anti-SDC1 (Thermofisher Scientific, cat. 10593-1-AP) and anti-HLA-G (Thermofisher Scientific, cat. MA1-19219) immunostaining, respectively.

For STB differentiation, 20k iTSC were plated on 2.5 µg/ml Collagen IV (Corning, cat. 354233) pre-coated 96-well plate in TS medium one day before. The next day, media was changed into STB medium, composed of DMEM/F12 with GlutaMAX, 0.1 mM β-mercaptoethanol, 0.3% bovine serum albumin, 1% ITS-X, 4% knockout serum albumin, 2 µM forskolin, 2.5 µM Y-27632, and 1x Antibiotic-Antinycotic^14^. We kept the culture in normoxia and changed medium every 2 days. Immunocytochemistry (ICC) was performed by anti-SDC1 (Thermofisher Scientific, cat. 10593-1-AP) along with DAPI after 96 h differentiation.

For EVT differentiation, 30-50k iTSCs were resuspended in 30 µl cold-Matrigel (Corning, cat. 354230) in a low attachment microtube and plated as a droplet on 1 µg/ml Collagen IV pre-coated 24-well plate. After allowing Matrigel solidification in a 37°C incubator for 15 min, EVT 1 medium was added on top of the droplet for 3 days. EVT II medium was added for 3 days replacing EVT 1 and finally EVT III medium was added for another 3 days replacing EVT II. ICC was performed with anti-HLA-G (Thermofisher Scientific, cat. MA1-19219) and anti-KRT7 (Thermofisher Scientific, cat. 181598) along with DAPI. Three EVT media were composed as followed: EVTI (DMEM/F12 with GlutaMAX, 0.1 mM β-mercaptoethanol, 0.3% bovine serum albumin, 1% ITS-X, 1 x Antibiotic-Antimycotic, 4% KSR, 7.5 µM A83, 2.5 µM Y-27632, 100 ng/ml NRG1 and 2% Matrigel), EVTII (DMEM/F12 with GlutaMAX, 0.1 mM β-mercaptoethanol, 0.3% bovine serum albumin, 1% ITS-X, 1 x Antibiotic-Antimycotic, 4% KSR, 7.5 µM A83, 2.5 µM Y-27632, and 0.5 % Matrigel), and EVTIII (DMEM/F12 with GlutaMAX, 0.1 mM β-mercaptoethanol, 0.3% bovine serum albumin, 1% ITS-X, 1 x Antibiotic-Antimycotic, 7.5 µM A83, 2.5 µM Y-27632, and 0.5% Matrigel).

### Cell lines

CT29 and CT30 were described in Okae et al 2018^14^. CT29 was obtained from Riken, Japan; CT30 was kindly provided by Dr. Theunissen. SCTi003-A and SCTi006-A hiPSC lines were procured from StemCell Technologies (cat 200-0511 and 200-0945) and cultured on 1% Matrigel-coated plates (Corning, cat 356230) in mTeSR Plus medium (StemCell Technologies, cat 100-0276) for three passages before processing for RT-qPCR.

### RNAseq

RNA was extracted from iTSCs cultured under normoxia (n = 23) and hypoxia (n = 23) using standard procedures. Libraries were prepared with the TruSeq Stranded Total RNA with Ribo-Zero H/M/R Gold kit and sequenced on a NovaSeq 10X Plus platform. RNA-seq data were processed using the SPEAQeasy pipeline^59^. Differential expression analysis was conducted using the limma package (v3.66.0)^60^, after filtering out lowly expressed genes with the filterByExpr function and adjusting for diagnosis, sex, RNA Integrity Number (RIN), and overall mapping rate. GO enrichment analysis of the significant DEGs (FDR < 0.05) was performed using enrichGO with default parameter settings.

### Cell Painting assay

Cells grown as described above were seeded in ViewPlate black 96 well plates (Revvity) at a seeding density of 40,000 cells/well. For select lines where attachment was poor (1002.01 and 1013.04) seeding density was increased so that density during imaging matched the remaining lines. Within a single differentiation, each cell line was seeded in four separate wells (n=4) across two plates. All cell lines underwent three independent differentiations (n=3) for a total of n=12 replicates per cell line per condition (i.e., normoxia and hypoxia). Cell painting was carried out under the standard protocol as described in the updated Bray et al protocol by Cimini et al^61^ using the PhenoVue Cell Painting kit (Revvity). Briefly, 50 µL of PhenoVue 641 mitochondrial staining solution was added to each well containing 100 µL of growth medium. Cells were incubated for 30 min at 37 °C. Cells were fixed by adding 50 µL of 16% (wt/vol) methanol-free PFA (final concentration 4% wt/vol) and incubating for 20 min at RT. Wells were washed 4x with 200 µL of 1x HBSS and aspirated. Staining solution containing the remaining PhenoVue reagents was added to each well (50 µL), and plates were incubated at RT for 30 min. Wells were washed 4x with 200 µL of 1x HBSS followed by the addition of 200 µL 1x HBSS to each well to prepare for imaging.

Plates were imaged on an ImageXpress Micro XLS widefield high-content imager (Molecular Devices). Imaging was carried out on 16 sites/well using a 20x/0.45 PanFluor objective, with DAPI, GFP, TRITC, TexasRed, and Cy5 channel filter sets used to visualize DNA, endoplasmic reticulum (ER), RNA/nucleoli, actin/Golgi/periplasm (AGP), and mitochondria, respectively.

### RT-qPCR

Total RNA was extracted from hiPSC lines using TRIzol Reagent (Ambion) and treated with DNase I (Zymo Research) to remove contaminating DNA. DNase-treated RNA (600 ng) was reverse-transcribed using the SuperScript IV First-Strand Synthesis System (Invitrogen). qPCR was performed using the QuantiTect SYBR Green PCR Kit (Qiagen) on the QuantStudio 3 Real-Time PCR System (Applied Biosystems). Reactions were conducted in technical triplicates and normalized to GAPDH. Relative gene expression was calculated using the 2^-ΔΔCT^ method normalized to female control line SCTi003-A (StemCell Technologies, cat 200-0511). Male control line SCTi006-A (StemCell Technologies, cat 200-0945) was used as a negative control for XIST expression. Primer sequences were validated in Raposo et. al, 2025^62^: XIST Fw: ACATGCCTGGCACTCTAGCA, XIST Rv: AAACATGGAAATGGGTAAGACACA, GAPDH Fw: GTCGTGGAGTCCACTGGCGTC, GAPDH Rv: TCATGAGTCCTTCCACGATAC.

### Image processing and feature extraction

Images were analyzed separately by (1) CellProfiler, (2) a pre-trained CNN developed by Spring Science, and (3) the DINOv3 Vision Transformer. (1) For CellProfiler analysis, images were first illumination corrected before performing feature extraction, using CellProfiler pipelines based on the JUMP_illum_LoadData_v1 and JUMP_analysis_v3 pipelines made available by the Broad Institute (https://github.com/broadinstitute/imaging-platform-pipelines/tree/master/JUMP_production). Analysis was performed on the Joint High Performance Computing Exchange (JHPCE) high-performance computing cluster at the Johns Hopkins Bloomberg School of Public Health. 3,672 features/cell were collected in this manner. CellProfiler output was analyzed using the Pycytominer python package from the Cytomining software ecosystem (pipeline adapted from https://github.com/cytomining/pipeline-examples/blob/main/)^63^. Briefly, single cell data from a single plate was compiled and median aggregated to the well level. Well level profiles were annotated with relevant metadata (e.g., sex and diagnosis), followed by whole-plate normalization by MAD robustization. Single plate profiles were combined, and feature selection was performed to remove redundant and highly correlated features, resulting in the profile used for downstream analysis. Feature selection resulted in well-level profiles of 486 features/well. An additional 46 features correlated to plate ID were removed following manual inspection, resulting in profiles of 440 features/well. For quality control, focus score was calculated by Spring Science. Focus score is a measure of the intensity variance across the image. This score is calculated using a normalized variance, which was the best-ranking algorithm for brightfield, phase contrast, and DIC images. Higher focus scores correspond to lower blurriness. (2) For CNN analysis, cell nuclei were identified by CellPose^64^. An 82px x 82px crop around the centroid of each nucleus was generated for each channel as input. The pre-trained CNN developed by Spring Science is proprietary and utilizes the RepLKnet CNN architecture^36^. 128 embeddings were extracted for each channel, making the total number of embeddings extracted 640. Resulting single cell profiles were median-aggregated to the per-site level and median aggregated again to the per-well level. (3) Features were extracted using a standard DINOv3 small (21m param) model. Nuclei were again identified by CellPose. For each image, patches of 80px x 80px were cropped around the centroid of each nucleus. Patches were resized to 224px x 224px before being fed into DINOv3. Per image standard normalization was applied using image-level statistics within each patch. These patches were fed into the model one channel at a time, with channels replicated 3 times to fit the required 3 channel input shape of DINOv3. The small DINOv3 produced 384 dimensional features per single cell at this stage. Site-level features were generated by taking the mean of all single cell features. Well-level features were generated by taking the mean of all site-level features.

### Feature profile analysis

*Hierarchical clustering*. Similarity matrices of well-level data were constructed using Pearson correlation coefficients followed by hierarchical clustering of the Pearson correlation coefficients using Morpheus by The Broad Institute (RRID:SCR_017386). All other analyses were performed in RStudio (Posit team (2025). RStudio: Integrated Development Environment for R. Posit Software, PBC, Boston, MA. URL http://www.posit.co/). *UMAPs*. UMAPs were performed using the umap R package with default parameters. *Machine learning classifiers*. Random Forest (RF), logistic regression (LR), and multilayer perceptron (MLP) classifiers were performed using the randomForest, tidymodels, and neuralnet R packages, respectively. For all RF classifiers, a ntree = 2500 was used along with the default mtry. All LR classifiers were run using the glmnet model with mixture = 0 (ridge regression) and penalty = 0. All MLP classifiers were run using two hidden layers of five and three units. Model performance was measured by area under the curve (AUC) of the receiver-operator characteristic (ROC) curve using the pROC R package. All classification tasks were performed using 10-fold cross-validation, where each fold contained two to three cell lines, with at least one positive and negative case (e.g. SCZ vs Ctrl). In each run, one fold was held out for subsequent testing, while the remaining nine folds were used to train the model. Performance was recorded as the median ROC AUC value across 10 folds. Feature importance lists generated during RF modeling were used to rank the most important features for sex and diagnosis classification tasks.

### Statistics

Quality control metrics (e.g., focus score and cell count) were analyzed by a Welch’s ANOVA, assuming a Gaussian distribution of residuals and unequal standard deviations. AUC ROCs and CellProfiler feature distributions were analyzed by a Mann-Whitney U test as, in general, we cannot assume a Gaussian distribution of AUCs or feature values. P values of < 0.05 were considered significant.

## Acknowledgements

We are grateful to the Lieber and Maltz families for their visionary support that funded part of this project. Tomoyo Sawada, Yanhong Wang, Bruno Araujo and Jennifer Erwin from the LIBD iPSC core, and Srinidhi Sripathy from LIBD for reprogramming and characterizing the 23 iPSC lines used in this study. The collection of skin fibroblasts was supported by direct funding from the Intramural Research Program (IRP) of the NIMH to the Clinical Brain Disorders Branch (DRW, PI, Protocol 95-M-0150, NCT00001486, Annual Report no. ZIA MH002942053) with supplemental support from the Clinical and Translational Neuroscience Branch (Karen Berman, PI). We thank Karen Berman, all of the participants in the IRP study and their families, and the families of the brain donors. The JHUSOM Microscopy Facility for assistance with the IXM high-content imager. Maggie Vantagoli Policelli and Aaron Waterman from Spring Science for providing guidance and consultations regarding the early implementation of CNN. Anne Carpenter from the Broad Institute for advice on Cell Painting best practices. James Barrow for critical reading of the manuscript and for supporting the project since its inception. Ingrid Buchler, Michael Poslusney, Gregory Carr from the Drug Discovery team, and Giovanna Punzi and Thorold Theunissen for helpful discussions. Joo Heon Shin for supervising the generation of RNAseq data. The Society for Biomolecular Imaging and Informatics (SBI2, https://sbi2.org/) for providing an invaluable venue to discuss state-of-the art methods and analytical tools. This work was supported by the Maryland Stem Cell Research Foundation Launch Grant 2024-MSCRFL-6392 (GU), Grant Number 5T15LM007359 from the National Library of Medicine (JP), and LIBD core fund (ES and GU). JHSOM Microscope Facility; this work was supported by the National Institutes of General Medical Sciences under award numbers R01GM28007-S1 and R01GM66817-S1. Figure 1a created with BioRender (https://biorender.com/), licensed under CC BY 4.0.

## Author contributions

GU, JK, DRW and ES conceived the study. FP, JK, GU, DRW and ES designed the study. FP and JK generated data. JP and JC performed DINOv3 analysis. JH processed and analyzed the RNAseq data. BS contributed to the culturing and imaging of iTSC. JJ contributed to the development of the Cell Painting pipeline and performed XIST analysis. TMH oversaw the donation, consenting process and clinical curation of the primary dural fibroblasts. BJM oversaw the reprogramming of skin fibroblasts to hiPSCs, curation and quality control of hIPSCs. FP and ES analyzed the data and wrote the first draft of the manuscript. FP, GU, JK, and ES wrote the final manuscript, which was revised and approved by all the authors

## Data Availability

Data is available upon request

## Code availability

All code used in this study is located at https://github.com/LieberInstitute/tsc-cellpainting/

## Supplemental Figures

**Supplemental Figure 1:**
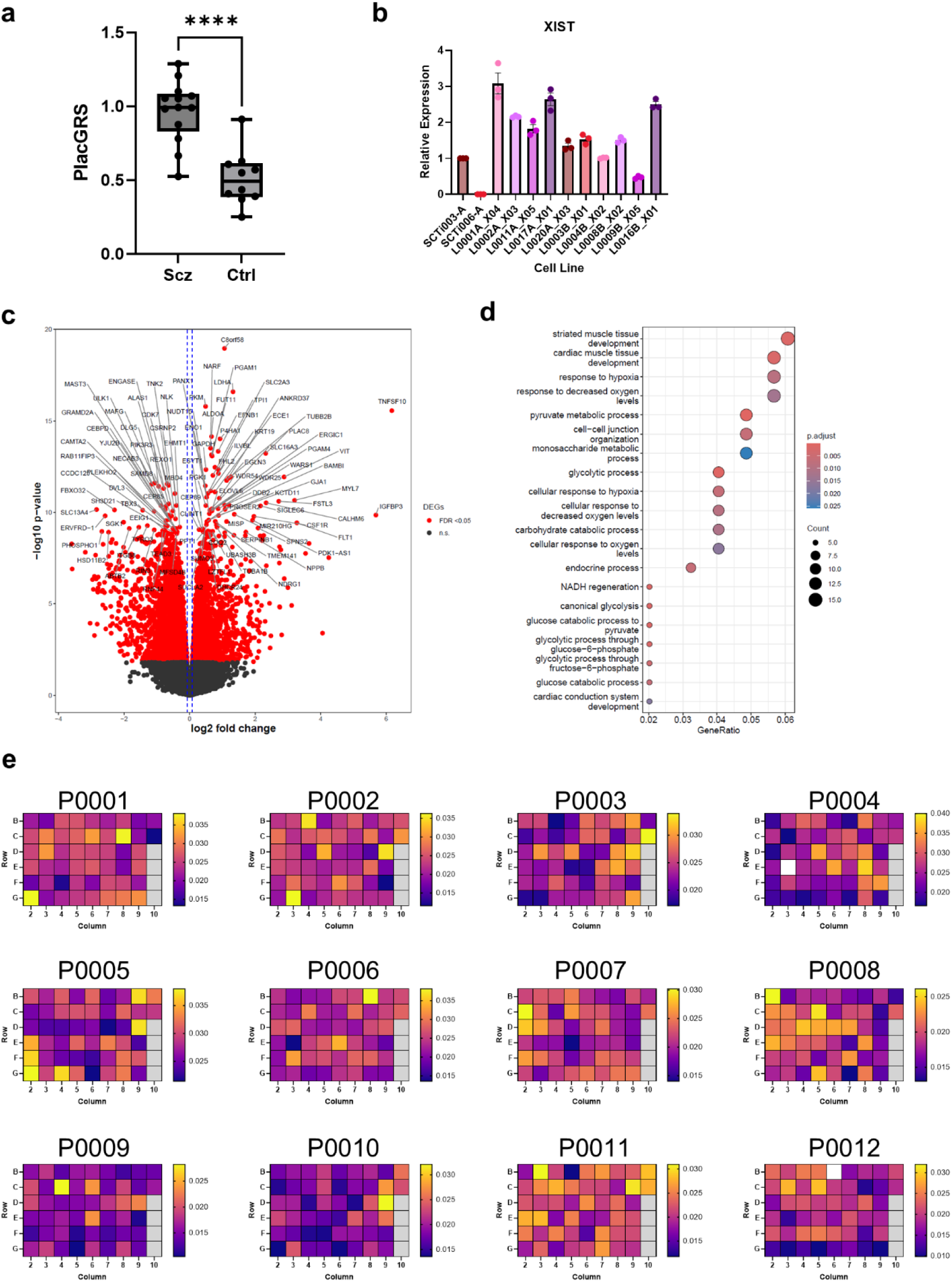
**Additional quality control of iTSC line identity, hypoxia, and Cell Painting performance**. **a**) Welch’s t-test comparison of PlacGRS between Ctrl and SCZ groups. Box plots display the interquartile range with a line at the median value. Whiskers extend from minimum to maximum values (****: p < 0.0001). **b**) RT-qPCR analysis of *XIST* expression normalized to GAPDH in female hiPSC lines, compared to SCTi003-A. Male line SCTi006-A serves as a negative control for *XIST* expression. c) Volcano plot of the differentially expressed genes (DEGs) associated with hypoxia, with log2 fold change on the x-axis (positive values indicating genes upregulated in hypoxia) and log10 p-value on the y-axis. Red dots indicate genes significantly associated with hypoxia (FDR-adjusted p-value < 0.05); the top DEGs are labeled with gene symbols. d) Dot plot of Gene Ontology (GO) terms significantly enriched among the DEGs (FDR < 0.05). Dot color represents the FDR-adjusted p-value for enrichment, while dot size reflects the number of DEGs assigned to each GO term. **e**) Well-level focus scores (per-site median, DNA channel) by plate position for individual plates imaged in this study.

**Supplemental Figure 2:**
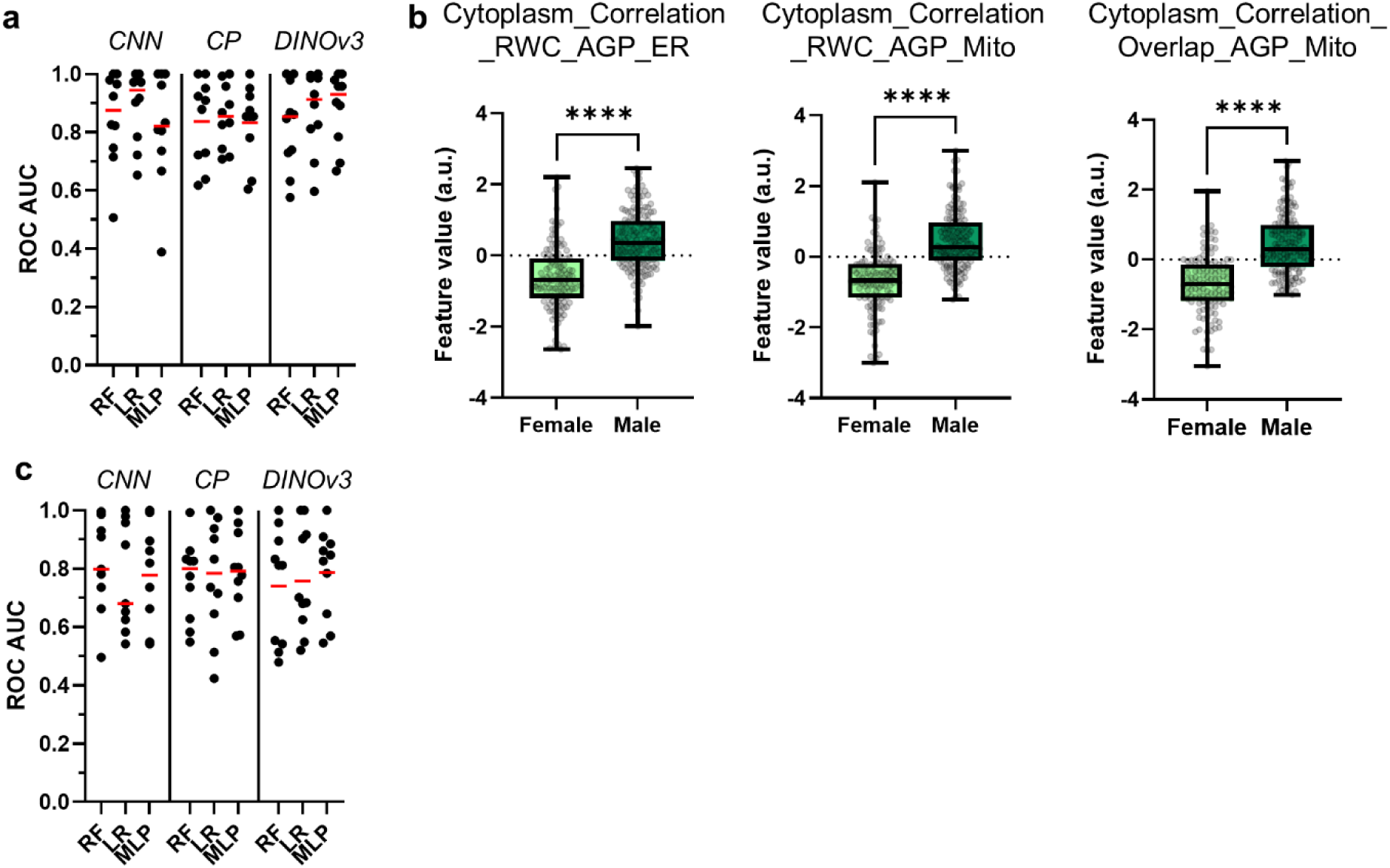
**Additional supervised machine learning model performance and feature importance under normoxia**. **a**) AUC ROC values for 10-fold cross validation of sex classification using CNN, CellProfiler (CP), and DINOv3 inputs. RF, LR, and MLP supervised learning models were applied to each dataset. Red line represents median value. **b**) Well-level comparison between male and female donor groups of *Cytoplasm_Correlation* features identified as highly important in RF sex classification modeling. Box plots display the interquartile range with a line at the median value. Whiskers extend from minimum to maximum values. All statistical analyses performed using a Mann-Whitney U test (***: p < 0.0001). **c**) Same as (a) but for diagnosis classification.

**Supplemental Figure 3:**
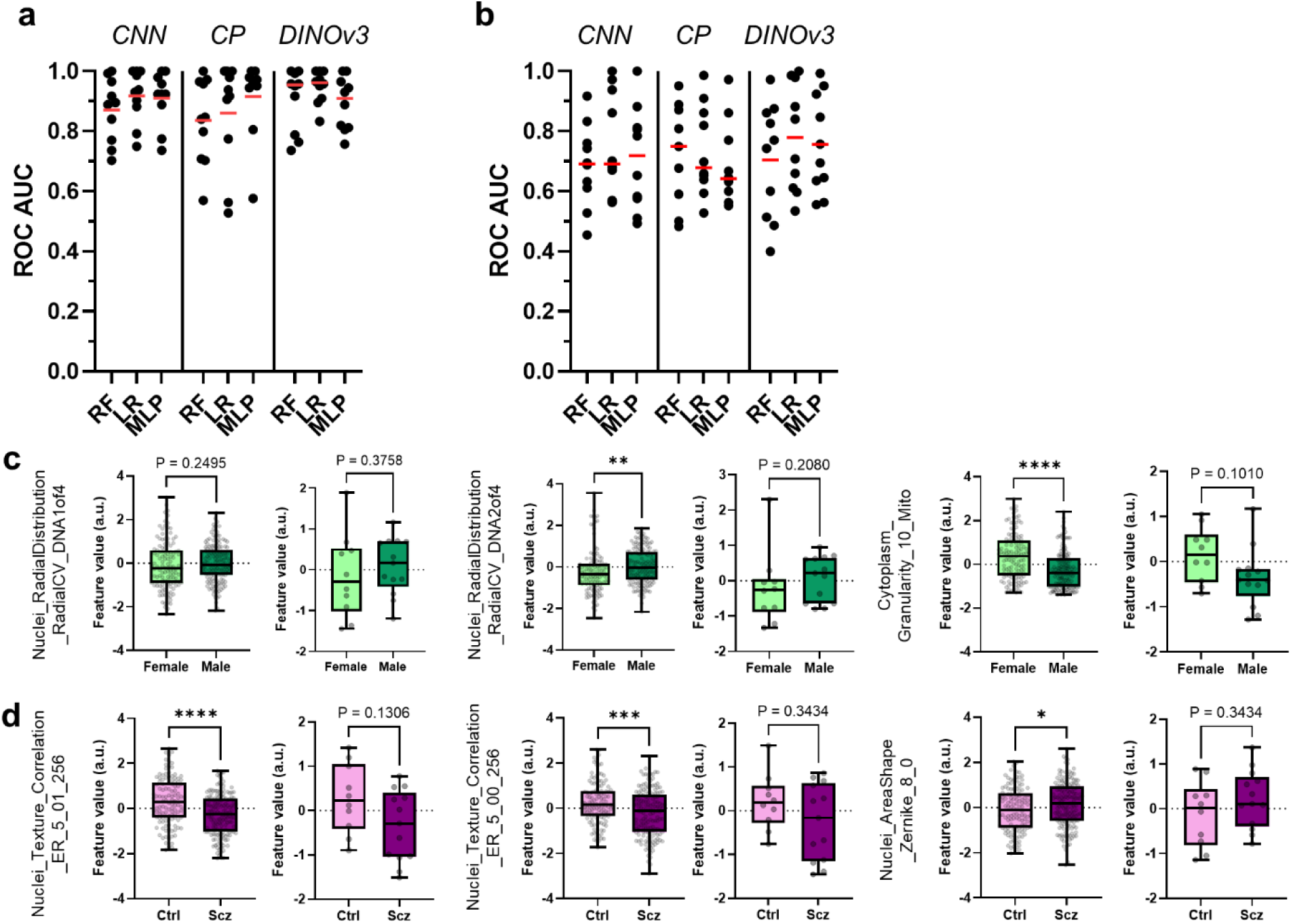
**Additional supervised machine learning model performance and feature importance under hypoxia**. **a**) AUC ROC values for 10-fold cross validation of sex classification using CNN, CP, and DINOv3 inputs. RF, LR, and MLP supervised learning models were applied to each dataset. Red line represents median value. **b**) Same as (a) but for diagnosis classification. **c**) Well (left) and cell line-level (right) comparisons between male and female donor groups under hypoxia for additional features observed as highly important for sex classification under normoxic growth. **d**) Same as (c) but for diagnosis classification. Box plots display the interquartile range with a line at the median value. Whiskers extend from minimum to maximum values. All statistical analyses performed using a Mann-Whitney U test (***: p < 0.0001).

**Supplemental Figure 4:**
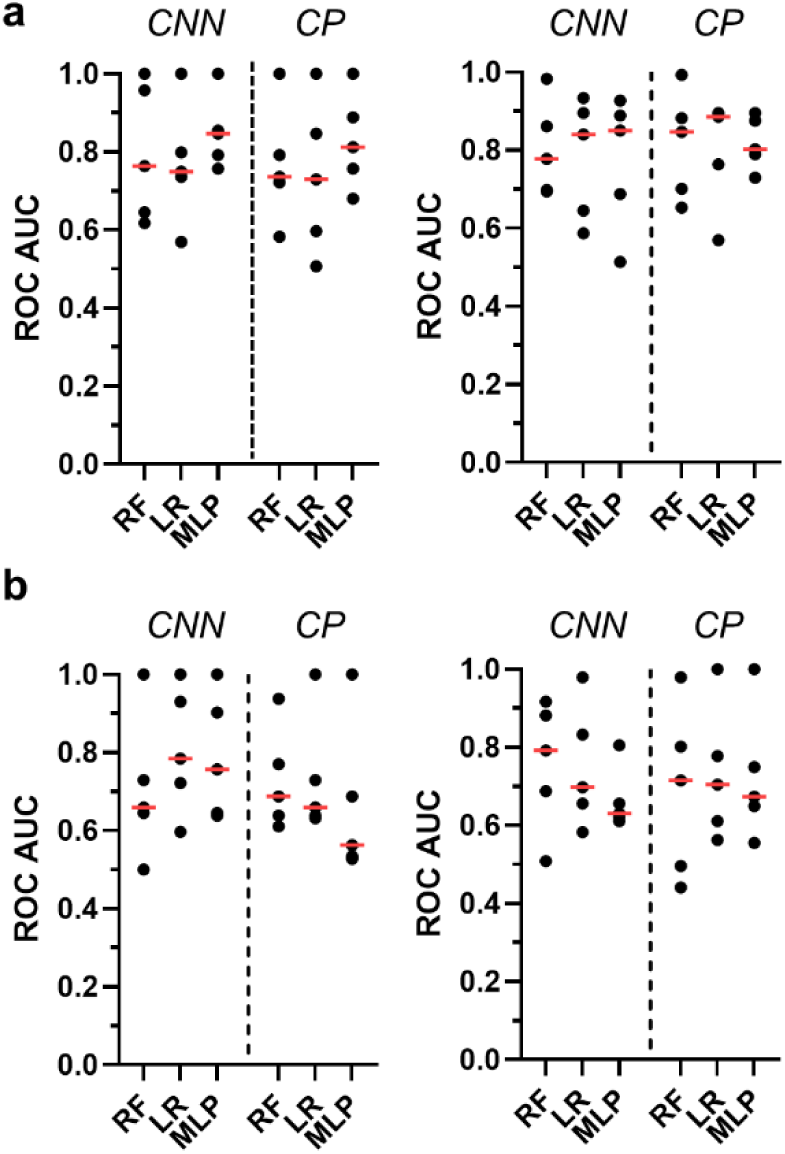
**Supervised learning model performance when trained and tested on exclusively male or female cell lines**. **a**) AUC ROC values for 10-fold cross validation of diagnosis classification using CNN and CP inputs. RF, LR, and MLP supervised learning models were applied to each dataset. Models were tested and trained exclusively on female (left) or male (right) cell lines grown under normoxia. **b**) Same as (a), but with models tested and trained exclusively on female (left) or male (right) cell lines grown under hypoxia. Red line represents median value.

## Supplemental Tables

Supplemental Table 1. Fold compositions for performing 10-fold cross validation of sex and diagnosis classifications (see Figures 3 and 4).

Supplemental Table 2. Importance values by fold from Random Forest modeling of sex classification task (see Figure 3).

Supplemental Table 3. Importance values by fold from Random Forest modeling of diagnosis classification task (see Figure 4).

